# Maize brace root mechanics vary by whorl, genotype, and reproductive stage

**DOI:** 10.1101/547794

**Authors:** Ashley N. Hostetler, Lindsay Erndwein, Elahe Ganji, Jonathan W. Reneau, Megan L. Killian, Erin E. Sparks

## Abstract

Root lodging is responsible for significant crop losses world-wide. During root lodging, roots fail by breaking, buckling, or pulling out of the ground. In maize, above-ground roots, called brace roots, have been shown to reduce root lodging susceptibility. However, the underlying structural-functional properties of brace roots that prevent root lodging are poorly defined. In this study, we quantified structural mechanical properties, geometry, and bending moduli for brace roots from different whorls, genotypes, and reproductive stages. Using 3-point bend tests, we show that brace root mechanics are variable by whorl, genotype, and reproductive stage. Generally, we find that within each genotype and reproductive stage, the brace roots from the whorl closest to the ground had higher structural mechanical properties and a lower bending modulus than brace roots from the second whorl. There was additional variation between genotypes and reproductive stages. Specifically, genotypes with higher structural mechanical properties also had a higher bending modulus, and senesced brace roots had lower structural mechanical properties than hydrated brace roots. Collectively these results highlight the importance of considering whorl-of-origin, genotype, and reproductive stage for quantification of brace root mechanics, which is important for mitigating crop loss due to root mechanical failure.

## INTRODUCTION

Roots are critical for plant health and productivity, including their pivotal role to anchor plants in the ground. In agriculture, a failure of plant anchorage causes significant crop loss and is referred to as root lodging (Carter and Hudelson 1988; Berry *et al.* 2004; Rajkumara 2008; Fedenko *et al.* 2015; Hostetler, Khangura, *et al.* 2021). During root lodging, roots fail by breaking, buckling, and/or pulling out of the ground (Easson *et al.* 1992; Ennos *et al.* 1993; Crook and Ennos 1993, 1994; Erndwein *et al.* 2020). With root breaking and buckling specifically (**Figure 1A**), roots fail when the mechanical load exceeds root structural tolerance. Despite the apparent role of root mechanics to limit root lodging, a detailed survey of root structural mechanical variation has not been performed (Stubbs *et al.* 2019).

**Figure 1.**
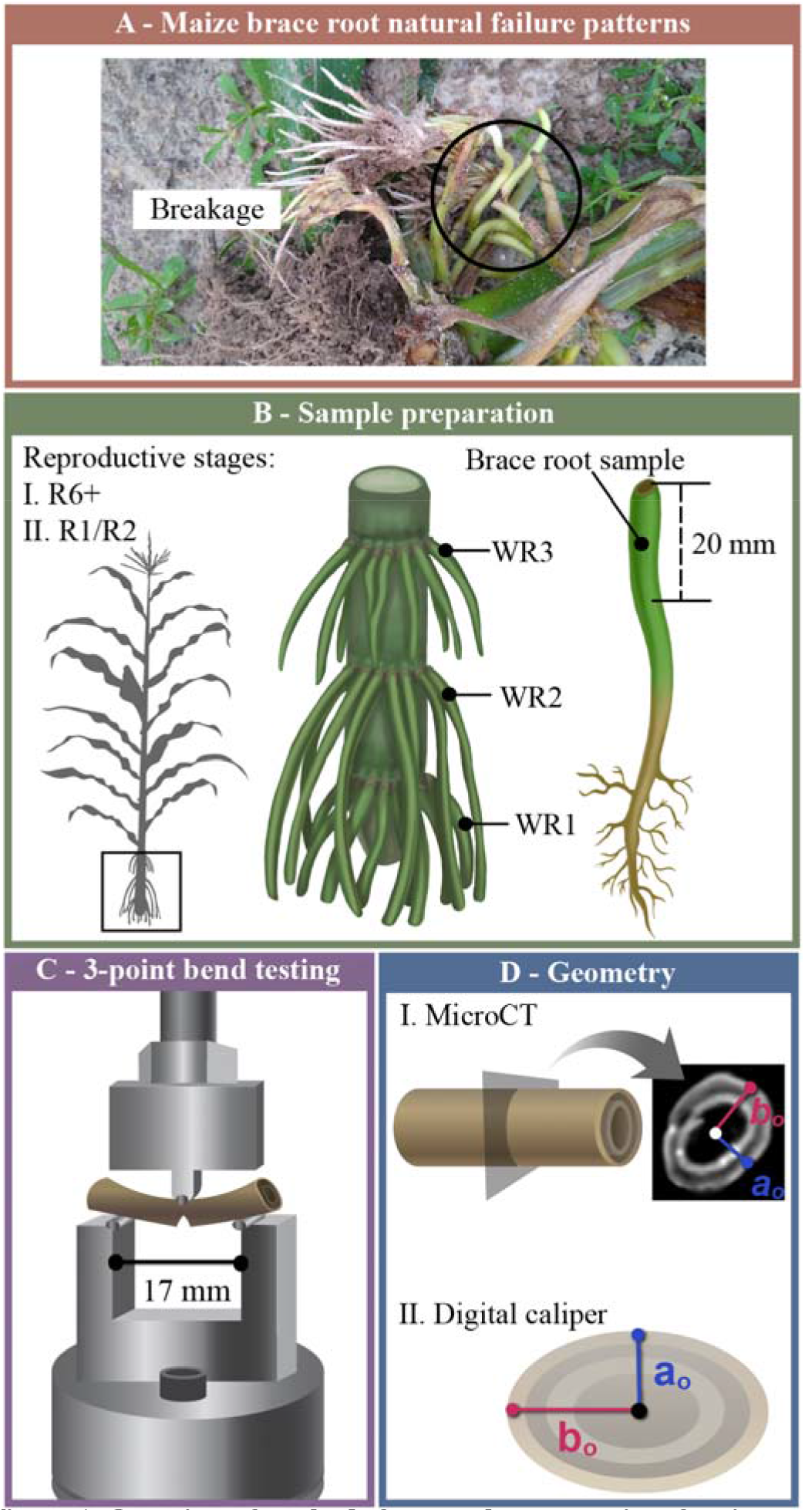
Overview of methods for sample preparation, 3-point bend testing, and quantification of brace root geometry. A) Maize brace roots break when the mechanical load exceeds tolerance. This failure is one mechanism of root lodging in maize. B) Maize brace roots were studied at the R6+ (senesced) and R1/R2 (hydrated) reproductive stages. Brace roots were collected from the whorl closest to the ground (WR1) and the second whorl (WR2). The first 20 mm of each brace root (closest to the stalk) was used for testing. C) The 3-point bend fixture was 17-mm with the load cell anvil applied to the center of the brace root sample at a constant rate of displacement until failure. Root failure is illustrated with the notch in the bottom of the brace root. D) Brace root geometry was quantified by either microCT scanning (R6+ samples) or digital calipers (R1/R2 samples).

In maize (*Zea mays*), root lodging causes between 7-25% yield losses in the United States, with the detrimental impact on yield increasing as plants reach reproductive maturity (Carter and Hudelson 1988; Tirado *et al.* 2021). The mature maize root system is composed of root whorls that develop from stem nodes both below (crown roots) and above (brace roots) the ground, which are collectively called nodal roots (Blizard and Sparks 2020). Previous studies have analyzed the structural mechanics of maize nodal roots via 3-point bending and shown that root mechanics are whorl-specific (Ennos *et al.* 1993; Goodman and Ennos 2001). These studies demonstrate that brace roots have higher structural mechanical properties and a lower bending modulus compared to crown roots (Ennos *et al.* 1993; Goodman and Ennos 2001).

In this study, we expand upon the previous studies and analyze the impact of whorl, genotype, and reproductive stage on maize brace root structural mechanics. We, and others, have shown that brace roots are critical for anchorage and root lodging resistance (Liu *et al.* 2012; Sharma and Carena 2016; Shi *et al.* 2019; Reneau *et al.* 2020; Hostetler, Erndwein, *et al.* 2021). We have further shown that there is variation in the brace root contribution to anchorage between whorls, with the whorl closest to the ground contributing the most (Reneau *et al.* 2020). Here, we address the hypothesis that brace root structural mechanics vary by whorl, genotype, and reproductive stage. To test this, we selected three temperate inbred lines (*Zea mays* L. cv. B73, Oh43, and A632) based on their agronomic importance (Liu *et al.* 2003) and variation in the brace root contribution to anchorage (Hostetler, Erndwein, *et al.* 2021). We subjected two whorls of brace roots to 3-point bend tests from these three inbred lines at two reproductive stages (hydrated and senesced brace roots). We show that the structural mechanical properties and bending moduli vary by genotype and reproductive stage. Together, these results highlight the importance of understanding brace root structural mechanics in the context of whorl-of-origin, genotype, and growth stage for future crop improvement.

## MATERIALS AND METHODS

### Plant material

Seeds from three inbred maize genotypes (*Zea mays* L. cv. B73, cv. Oh43, and cv. A632) were grown in the summers of 2019 and 2020. Seeds were planted on 05/22/2019 and 06/01/2020 in two replicate plots in Newark, DE (39° 40’ N and 75° 45’ W). Weather data for the Newark, DE field site can be found by selecting the Newark, DE-Ag Farm station on the Delaware Environmental Observing System (http://www.deos.udel.edu). In both years, fields were treated with a pre-emergence (Lexar at 3.5 quart per acre and Simazine at 1.2 quart per acre) and post-emergence (Accent at 0.67 ounces per acre) herbicide. At the time of planting, Ammonium Sulfate (21-0-0 at 90 lbs per acre, fertilizer) and COUNTER 20G insecticide (at 5.5 lbs per acre) were applied. Approximately 1-month after planting, when plants were at knee high, additional fertilizer was supplied (30% Urea Ammonium Nitrate at 40 gallons per acre).

In 2019, brace roots were collected at reproductive maturity/senescence (R6+, ~123 days after planting) from whorls that entered the ground, and designated as whorl 1 (bottom most whorl, whorl closest to ground), whorl 2 (subsequent whorl, whorl originating from second node above ground), and whorl 3 (whorl originating from the third node above the ground, **Figure 1B**). Brace roots were not collected if they showed signs of disease or splintered during collection. Brace roots were stored in coin envelopes until 3-point bend testing and micro-computed tomography (microCT) scanning. In 2020, brace roots were removed from whorl 1 and whorl 2 at the silking/blistering reproductive stage (R1/R2, ~67 days after planting). Brace roots collected at the R1/R2 stage were placed on a damp paper towel to maintain moisture and subjected to 3-point bend tests immediately following collection.

### Quantification of the brace root contribution to anchorage

The brace root contribution to anchorage (BRC) was determined *in situ* by comparing the slope of the Force-Deflection curve of maize plants with brace roots removed (None) to the slope of the Force-Deflection curve of maize plants with brace roots intact (All), as previously described (BRC=None/All; Reneau *et al.* 2020; Hostetler, Erndwein, *et al.* 2021). Plants were tested prior to the removal of brace roots and again after the removal of each subsequent whorl (starting with removal of the top-most whorl). These data enabled us to determine the overall contribution of brace roots to anchorage as well as the contribution of each individual whorl. A brace root contribution to anchorage ratio close to 1 indicates that anchorage was not impacted by the removal of brace roots, whereas a ratio close to 0 indicates that anchorage was predominantly dependent on brace roots entering the ground.

### Quantification of structural mechanical properties using three-point bending

A custom 3-point bend fixture was machined with a 17-mm span length (**Figure 1C**; **Supplementary Figure S1**), which was the longest span of a straight brace root section that could be reliably isolated. Brace roots were trimmed to include only the 20 mm of root closest to the stem (**Figure 1B**). At least two brace roots per whorl were tested for at least three plants per genotype and reproductive stage. Brace root samples were tested using an Instron 5943 (Norwood, Massachusetts USA) equipped with a 100 N load cell (Instron 2530 Series static load cell, Norwood, Massachusetts USA). Each brace root sample was placed on the fixture and adjusted for midpoint loading (**Supplementary Figure S1**). Prior to testing, each sample was preloaded to 0.2 N and 3-point bend tests were performed by constant rate displacement of the top fixture at 1 mm/min, until root failure. Force-displacement data were captured with Bluehill 3 software (Instron, Norwood, Massachusetts USA). Testing continued until failure, which was defined as the first steep decline in the force-displacement curve (**Figure 3A**) and characterized as a crack forming in the brace root sample opposite of the loading site. For complex biological tissues such as brace roots, the underlying cell layers likely have different structural mechanical properties that contribute to the overall tissue mechanics. For example, roots are often considered structured as two concentric hollow cylinders of lignified tissue (Ennos *et al.* 1993; Chimungu *et al.* 2015). However, the structural mechanical properties of individual brace root cell layers are unknown, thus we consider only the properties of the entire root in this study (Niklas and Spatz 2012). Structural mechanical properties were extracted from force-displacement curves (**Figure 3A**) using Bluehill 3 software. Structural stiffness (K) was defined as the linear slope of the force-displacement curve; ultimate load (UL) was defined as the maximum force the sample withstood before failure; break load (BL) was defined as the force upon fracture, illustrated as the sharp drop in the force-displacement curve (**Figure 3A**).

**Figure 2.**
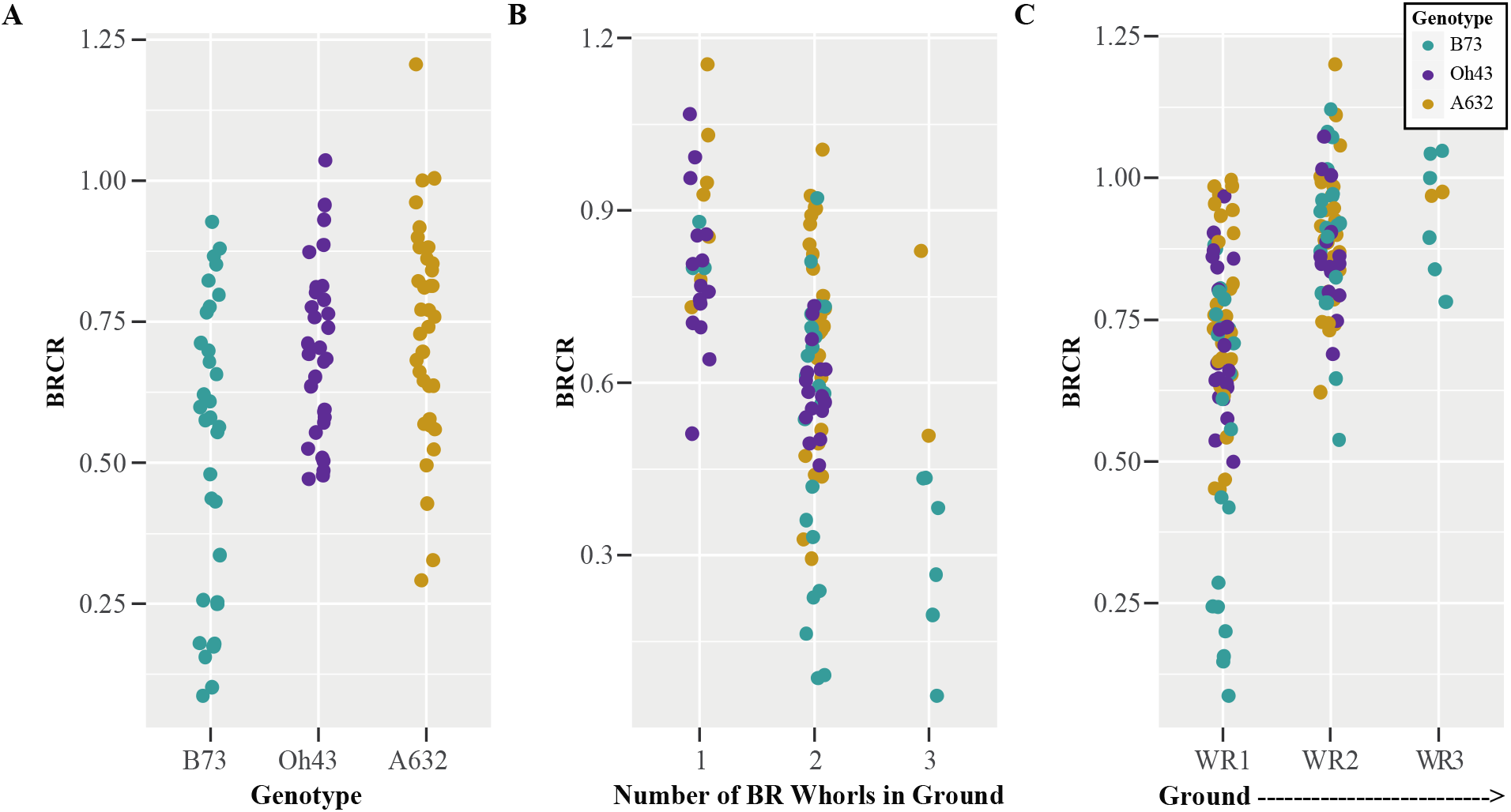
Brace roots are important for anchorage and whorls closer to the ground contribute more at the R6+ reproductive stage. A) The brace roots from three maize genotypes (B73, Oh43, and A632) had variable contributions to anchorage. B) The brace root contribution to anchorage was greater when there were more whorls of brace roots in the ground. C) The brace root whorl closest to the ground (WR1) contributed the most to anchorage. BR – brace root. WR – whorl. BRCR - <u>B</u>race <u>R</u>oot <u>C</u>ontribution <u>R</u>atio; BRCR close to 1 indicates anchorage is not impacted by the removal of brace roots, whereas a BRCR close to 0 indicates anchorage is dependent on the number of brace root whorls entering the ground.

**Figure 3.**
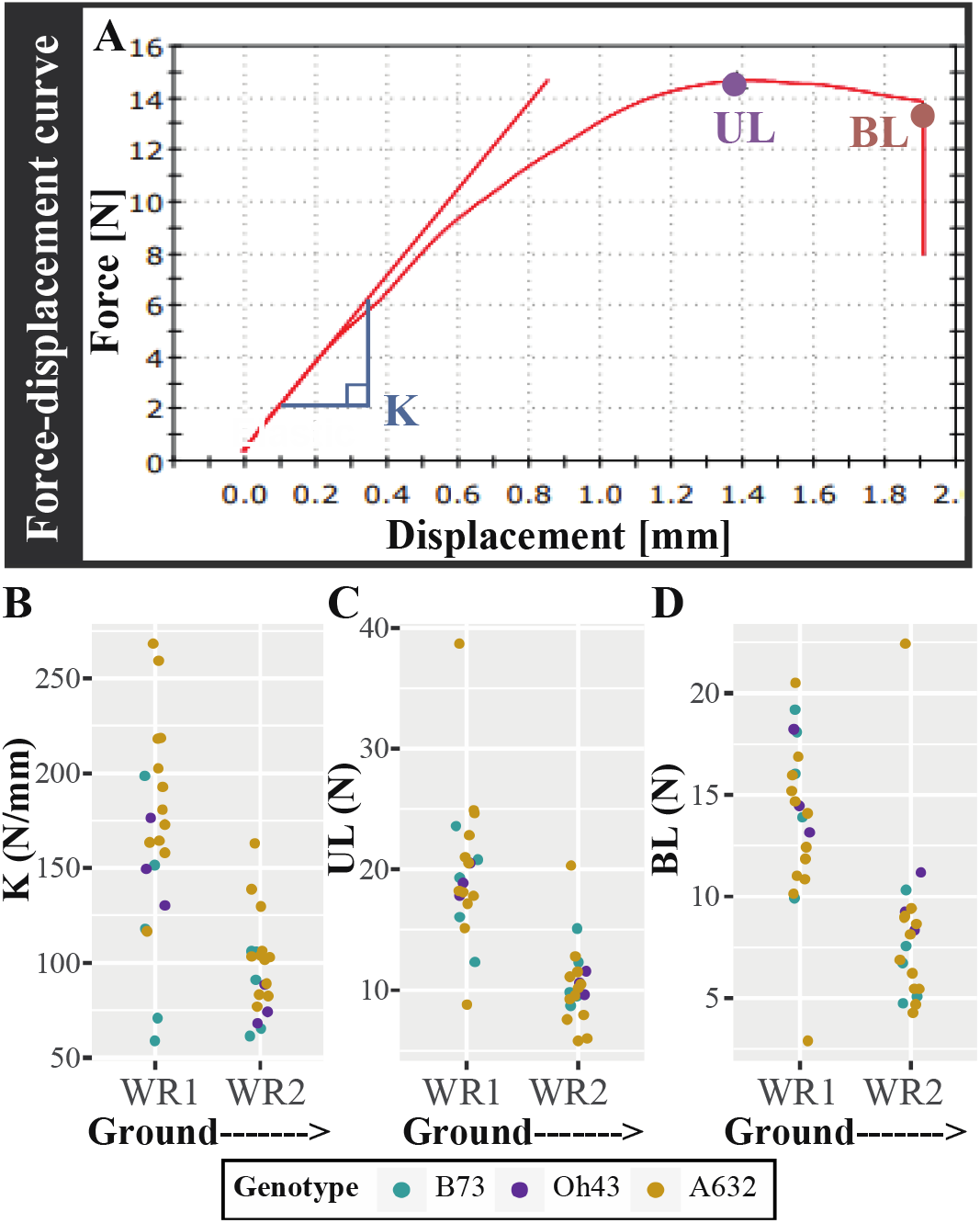
The structural mechanical properties of maize brace roots are greater for whorls closer to the ground at the R6+ reproductive stage. A) Example of a force-displacement curve from a R6+ brace root subjected to 3-point bending. B) An idealized force-displacement curve to illustrate the structural mechanical properties that were extracted. The structural stiffness (K) was defined as the linear slope of the force-displacement curve, the ultimate load (UL) was defined as the highest force the sample withstood without failure, and the break load (BL) was defined as the force upon fracture. C-E) A two-way ANOVA showed that: C) K was impacted by both whorl (p≤0.05) and genotype (p≤0.05), with whorls closer to the ground (WR1) having a higher K than whorls farther away (WR2), and A632 having higher K compared to B73. D) UL was impacted by whorl (p≤0.05), with whorls closer to the ground (WR1) having a larger UL. E) BL was impacted by whorl (p≤0.05), with whorls closer to the ground (WR1) having a larger BL. WR – whorl.

### Quantification of brace root geometry

Sample geometry was measured in two ways based on sample constraints. For the senesced root samples (R6+), brace root geometry was quantified by microCT (**Figure 1D**) analysis. For microCT analysis, brace root samples were inserted into a low density upholstery foam fixture to provide a supportive bed that would not appear on the scan (**Supplementary Figure S2**) and samples were scanned with a Bruker Skyscan 1276 (voxel size of 42.32 μm, 40 kV, 100 μA, 203 msec exposure time, angular step of 0.4 degrees). Scans were reconstructed using Bruker Nrecon software, and Fiji software (Schindelin *et al.* 2012) was used to measure the brace root radii from the center of microCT scans to the exterior of the sample (ao - the minor axis perpendicular to bending and b_o_ - the major axis parallel to bending) (**Supplementary Figure S2D**). Brace root diameter was determined by doubling radii (a_o_ and b_o_) quantifications.

In contrast, fresh tissue (R1/R2) geometry was measured immediately upon collection with a digital caliper (DC) (NEIKO 01407A, 0-6 inch), which minimized the time to testing and brace root dehydration. Brace root samples are frequently ovular; therefore, the largest diameter (majorDC) and smallest diameter (minorDC) (**Figure 1D**) were measured at the midpoint of the brace root section, which is the loading site during 3-point bend testing (**Figure 1C**; **Supplementary Figure S1**). The sample radii (a_o_ - the minor axis perpendicular to bending and b_o_ - the major axis parallel to bending), were calculated by taking half of the digital caliper measurements.

### Quantification of the second moment of area (*I*)

Quantifying the area through which a stress operates is challenging for irregular biological structures such as brace roots (Niklas and Spatz 2012). Therefore, previous studies assessing root mechanics have used a geometric simplification and considered the root a solid cylinder (Ennos *et al.* 1993; Goodman and Ennos 1996, 1997, 1998, 2001). In this study, we have a high-resolution geometric quantification of the brace roots from microCT scans (**Supplementary Figure S2**), thus allowing us to evaluate this geometric simplification for calculating the second moment of area (*I).* The second moment of area of R6+ brace root samples were calculated using two different approaches 1) true, where the second moment of area is calculated directly by CT Analyzer software, and 2) simplified, where brace root geometry is considered a solid cylinder as in previous studies (Eq. 1):

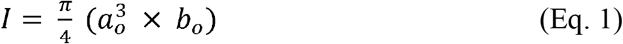

where π = 3.1415, a_o_ is the minor axis perpendicular to bending, and b_o_ is the major axis parallel to bending. The true second moment of area was calculated from the microCT scanned images. For each sample, images were resliced along the root axis (Data Viewer, Brunker, Belgium), and cross-sectional stacks were then thresholded to separate the brace roots from the background. The second moment of area was then calculated with the 2D analysis function for each thresholded sample at the mid-length, using CT Analyzer software (Bruker, Belgium). Three values of the true second moment of area were extracted from the CT Analyzer software: *I*_true_ major, *I*_true_ minor, and *I*_true_ average.

*I* and *I*_true_ were compared with a Pearson correlation analysis in R ver. 4.0.2 (R Core Team 2013). Comparison of the *I*_true_ (major, minor, and average) with the simplified *I* assumption showed high positive correlations (r > 0.83) (**Supplementary Figure S3**). Based on the high correlation between these data, we conclude that a simplified geometry to a solid cylinder, as used in previous studies (Ennos *et al.* 1993; Goodman and Ennos 1996, 1997, 1998, 2001), is a reasonable approximation for brace root geometry. Thus, the second moment of area of the brace roots was calculated with simplified geometry (Eq. 1), and *I* was used throughout the paper to enable comparison between this study and previous results.

### Quantification of material properties

The structural bending modulus (E) was calculated using the equation for a center-loaded beam supported at both ends by a lower fixture (Eq. 2 adapted from (Al-Zube *et al.* 2018)).

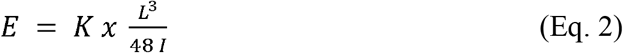

where K is the structural stiffness, L is the fixture span length (17 mm for samples tested here), and *I* is the second moment of area (calculated with simplified geometry, Eq. 1).

### Statistical analysis

All statistical analyses were performed in R ver. 4.0.2 (R Core Team 2013). Data used in this manuscript are as follows: contribution to anchorage (overall contribution; whorl 1 ratio; whorl 2 ratio; whorl 3 ratio), structural mechanical properties (K; UL; BL), brace root geometry (major diameter; minor diameter, and the second moment of area (*I*)), and material properties (bending modulus (E)). An analysis of variance (ANOVA) was used for all statistical tests. A Shapiro-Wilk test was used to determine if data were normally distributed, and when data were not normally distributed (p≤0.05), data were transformed with Tukey’s ladder of Powers from the *rcompanion* package ver. 2.3.26 (Mangiafico 2021). Transformed values were then used in the ANOVA. If a significant difference was found (p≤0.05), a post-hoc Tukey Honest Significant Difference (HSD) test was used to test all pairwise comparisons.

For contribution to anchorage data, a two-way ANOVA was run to test the effect of genotype (B73, Oh43, A632) and number of brace root whorls in the ground (1 whorl, 2 whorls, 3 whorls) on the overall contribution to anchorage, and the effect of genotype and individual whorls (whorl 1, whorl 2, whorl 3) on the individual whorl ratio. For structural mechanical properties, brace root geometry, and material properties, data from brace roots within the same whorl for each plant were averaged to provide a single value per whorl per plant. These data were then tested with a two-way ANOVA, where the effect of genotype (B73, Oh43, A632) and whorl (whorl 1, whorl 2) were considered within a reproductive stage (R1/R2 or R6+). Additionally, for structural mechanical properties and *I*, a three-way ANOVA was used to test the effect of genotype (B73, Oh43, A632), whorl (whorl 1, whorl 2), and reproductive stage (R1/R2, R6+). Pearson correlation analyses were run on structural mechanical properties and brace root geometry. All figures were generated in R with the *ggplot2* package ver. 3.3.3 (Wickham 2016).

### Data Availability

All raw data and R scripts used to process and analyze data are available at: https://github.com/EESparksLab/Hostetler_Erndwein_et_al_2021.

## RESULTS

### Brace root whorls closer to the ground have a greater contribution to anchorage

The inbred lines used in this study had a variable, but overlapping contribution of brace roots to anchorage (**Figure 2A, Tables S1-S2**; (Hostetler, Erndwein, *et al.* 2021)). The brace root contribution to anchorage is described by comparing the slope of the Force-Deflection curve with brace roots removed to the slope of the Force-Deflection curve with brace roots intact, with a ratio close to 1 indicating a low contribution of brace roots to anchorage and a ratio close to 0 indicating a high contribution of brace roots to anchorage (Reneau *et al.* 2020). For all genotypes, the brace root contribution to anchorage was greater when there are more whorls in the ground (**Figure 2B**, **Tables S1-S2**). Additionally, brace root whorls closer to the ground contributed more than brace roots from higher whorls for all three genotypes (**Figure 2C**, **Tables S1-S2**). These data confirm and expand upon our previous results (Reneau *et al.* 2020) to demonstrate that brace roots contribute to anchorage and the whorl closest to the ground has the greatest contribution relative to the other whorls. Since a third whorl (numbered in order of developmental progression from the ground up; **Figure 1B**) of brace roots is rarely observed in these genotypes (**Figure 2C**, **Table S2**, (Reneau *et al.* 2020)), we quantified the mechanics of brace roots from whorls 1 and 2 only (**Figure 1B**).

### Brace roots from whorls closer to the ground are more stiff

Brace roots were collected from the same plants that were measured for their brace root contribution to anchorage, and the structural mechanical properties of brace roots from whorl 1 and whorl 2 were measured using 3-point bend testing (R6+, **Figure 1B-C**, **Supplementary Figure S1**). For complex biological tissues such as brace roots, we consider the structural properties of the entire root (called structural mechanical properties throughout; (Niklas and Spatz 2012)). Consistent with a higher contribution to anchorage, structural mechanical properties were significantly higher for brace roots from whorl 1 compared to brace roots from whorl 2 regardless of genotype (**Figure 3B-D**, **Table 1, Tables S3-S4**). In addition to the differences between whorls within a genotype, there were also differences between genotypes for structural stiffness (K). Specifically, the K of A632 was higher than the K for B73 within each whorl, although genotypes did not differ for ultimate load (UL) or break load (BL) (**Tables S3-S4)**. These findings are consistent with the genotypic differences observed for the brace root contribution to anchorage (**Tables S1-S2**). Collectively, these results demonstrate that brace roots from whorls closer to the ground are stronger compared to brace roots from whorls further from the ground (higher on the stalk).

**Table 1.**
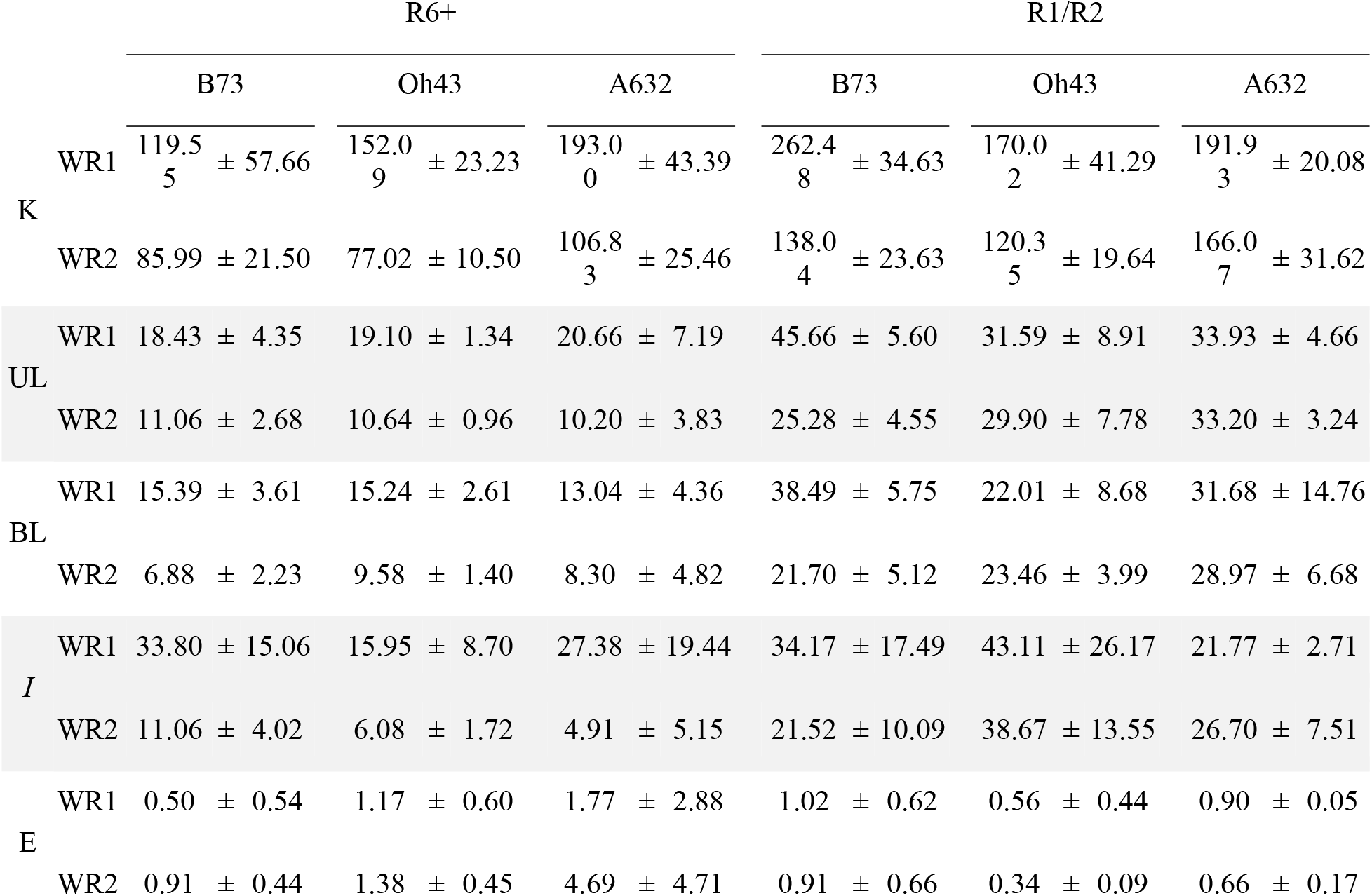
The impact of whorl, genotype, and reproductive stage on brace root structural mechanical properties, geometry, and bending moduli.

### Brace roots from whorls closer to the ground are larger

The differences in brace root structural mechanics between whorls may be related to the distribution of forces across different cross-sectional areas, where larger cross-sectional areas able to withstand greater forces. Previous studies have shown that nodal root diameters increase sequentially as root development progresses towards the shoot (Hoppe *et al.* 1986; Ennos *et al.* 1993), although these studies have primarily measured underground crown roots. Indeed, we found the opposite trend when considering just brace roots; both the major and minor diameters of brace roots were reduced as root development progressed from whorl 1 to whorl 2 (**Figure 4A-B**, **Tables S3-S4**).

**Figure 4.**
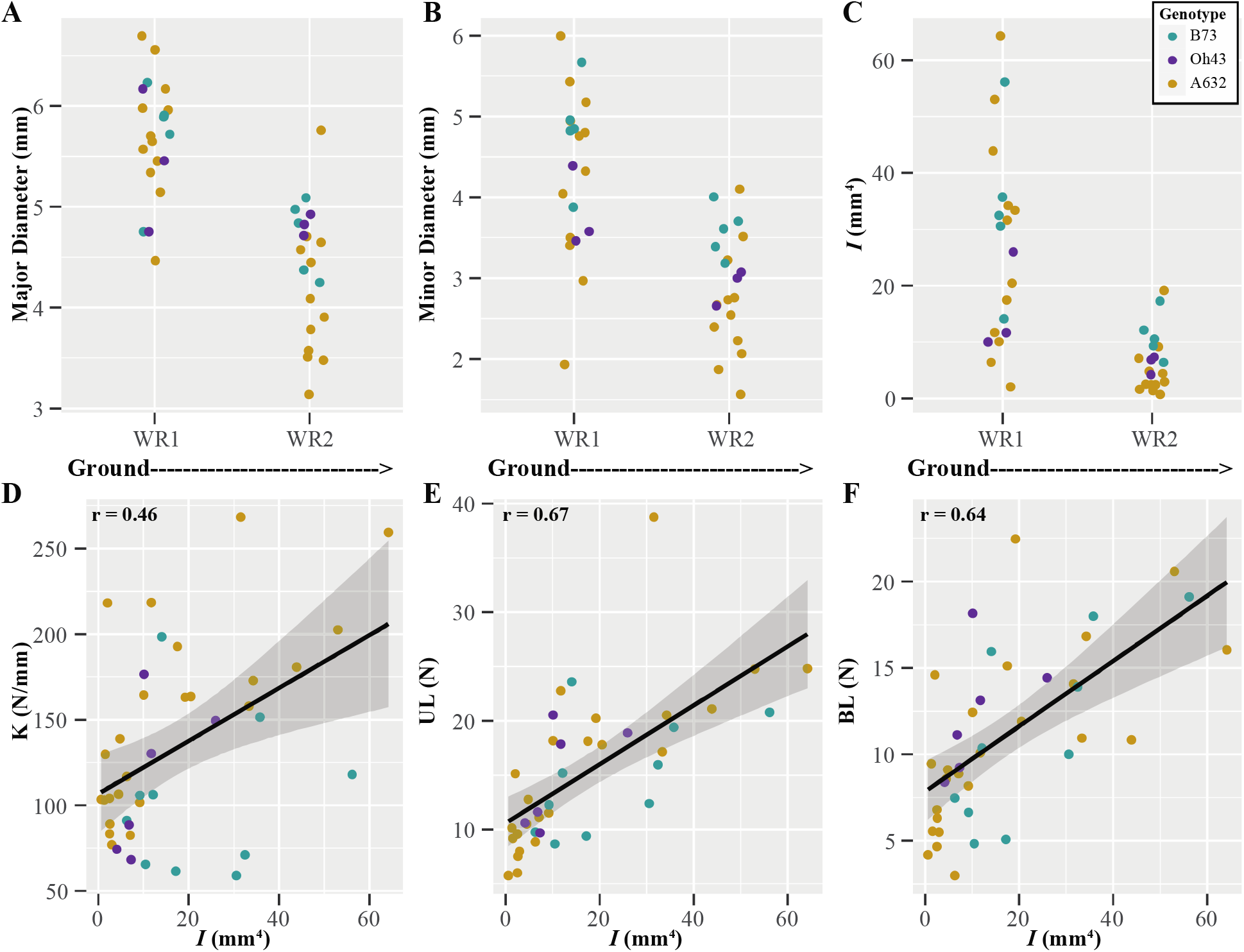
Brace root geometry partially explains structural mechanical properties with whorls closer to the ground being larger at the R6+ reproductive stage. A-C) A two-way ANOVA showed that: A) the major diameter of brace roots was impacted by whorl (p≤0.05), with whorls closer to the ground (WR1) having a larger diameter. B) the minor diameter of brace roots was impacted by both whorl (p≤0.05) and genotype (p≤0.05), with whorls closer to the ground (WR1) having a larger minor diameter, and B73 had a larger diameter than Oh43 and A632. C) the second moment of area (*I*) was impacted by both whorl (p≤0.05) and genotype (p≤0.05), with whorls closer to the ground (WR1) having a larger *I*, and B73 having a larger *I* than A632. D-F) Pearson correlation analyses showed that structural mechanical properties were positively correlated with the second moment of area (*I*). D) K was positively correlated with *I* (r=0.46). E) UL was positively correlated with *I* (r=0.67). F) BL was positively correlated with *I* (r=0.64). WR – whorl.

Additionally, the distribution of forces during bending can be described by the distribution of material away from the axis of the root - the second moment of area (*I*), which considers both the major and minor radii relative to the plane of bending. Like brace root diameter, *I* was larger for whorl 1 compared to whorl 2 (**Figure 4C**, **Table 1, Tables S3-S4**), and correlations between structural mechanical properties and *I* were positive and varied between r = 0.46 - 0.67 (**Figure 4D-F**). These results are consistent with larger roots having greater bending strength (**Figure 3**, **Figure 4, Tables S3-S4**). These results also highlight the importance of considering underground nodal roots (crown roots) (Hoppe *et al.* 1986; Ennos *et al.* 1993) separately from aboveground nodal roots (brace roots).

### The bending modulus of brace roots varies by whorl and genotype

Our results support a relationship between brace root geometry and structural mechanical properties; however, it is possible that there are also differences in the material properties of the brace roots (e.g., cell wall composition or structure). To assess material properties, the bending modulus (E) was calculated, which enables a universal comparison of the brace root’s ability to resist bending, with a higher value for E indicating a higher resistance to bending. Interestingly, brace roots from whorl 1 had lower E than brace roots from whorl 2 (**Figure 5**, **Table 1**, **Tables S3-S4**), which is opposite of what was observed for structural mechanical properties (**Figure 2**). In other words, whorl 2 has a higher resistance to bending (**Figure 5**), but an overall lower strength (**Figure 2**) compared to whorl 1 regardless of genotype. In addition, there was a genotype effect on E, with A632 having an overall higher E compared to B73, but neither genotype was different from Oh43 (**Tables S3-S4**). These genotypic relationships were the same relative to the relationships identified for K and *I* (**Figure 2-4**, **Tables S3-S4**). The difference in E between whorls and genotypes suggests an underlying difference in material properties between brace root whorls and genotypes. However, E was calculated from the geometric simplification of *I* as a solid cylinder. To ensure that this geometric simplification did not result in the differences observed between whorls, we measured the true second moment of area (*I*_true_) from microCT scans of each root, and calculated E from the *I*_true_. Calculation of E from *I*_true_ retained the differences between whorls and genotypes (**Table S5-S6**) indicating that the differences in E are due to underlying material properties, and not the geometric simplification.

**Figure 5.**
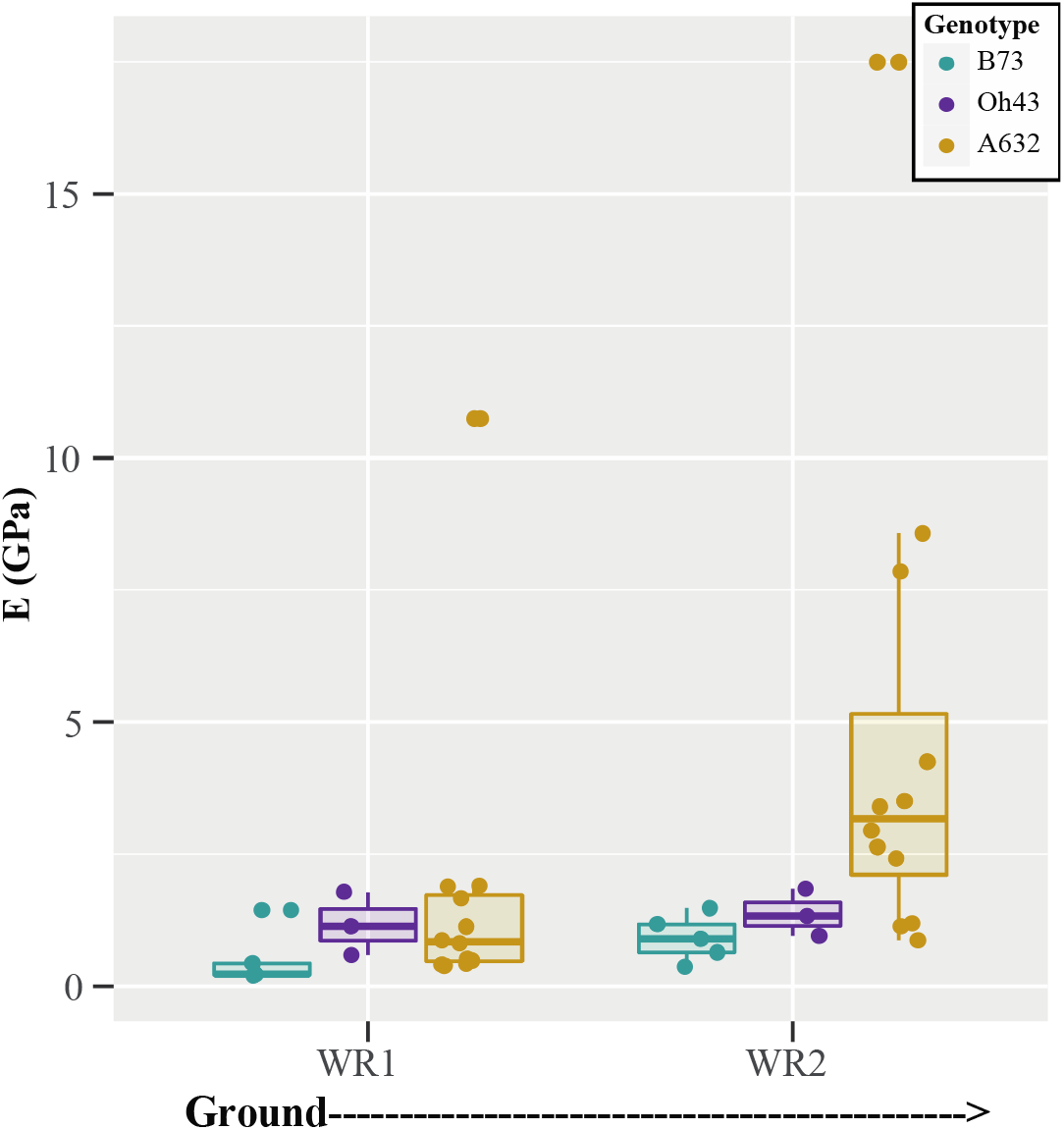
Brace root bending modulus varies by whorl and genotype at the R6+ reproductive stage. A two-way ANOVA sho3w4e9d that the bending modulus (E) of maize brace roots varied by both whorl (p≤0.05) and genotype (p≤0.05). The E of brace roots from whorls closer to the ground were lower than those from whorls higher on the stalk, and A632 had a higher bending modulus than B73. WR - whorl.

### Brace root structural mechanical properties and geometry are genotype-dependent at early reproductive stages

Brace root mechanics were originally assessed at the R6+ reproductive stage to mirror the timing of the assessment of the brace root contribution to anchorage. However, in 2020, a tropical storm caused root lodging at our Newark, DE field site (65 DAP; R1/R2 reproductive stage) and provided a unique opportunity to assess brace root mechanics at a growth stage when a natural lodging event occurred. To determine if results from senesced brace roots (R6+ stage) are consistent with hydrated brace roots (R1/R2 stage), we subjected R1/R2 reproductive stage brace roots from whorl 1 and whorl 2 to 3-point bend tests (**Figure 1C, Supplementary Figure S1**) and quantified brace root geometry.

Analysis of the structural mechanical properties at the R1/R2 reproductive stage showed that the effect of whorl was genotype-dependent (**Supplementary Figure S4**, **Tables S3-S4**). Specifically, B73 was the only genotype where hydrated brace roots mirror the results of senesced brace roots, with whorl 1 being stronger than whorl 2 for K, UL, and BL. Like B73, Oh43 had a higher K for whorl 1 compared to whorl 2, but there were no differences between whorls for the UL or BL. In contrast, A632 showed no differences between whorls for any of the structural mechanical properties. Unlike the senesced samples, the differences in structural mechanical properties were not explained by brace root geometry, with the geometries showing no significant difference by whorl and low correlations between structural mechanical properties and geometry (**Supplementary Figure S5; Tables S3-S4**). These results continue to refute previous conclusions that nodal root geometry increases with subsequent whorls (Hoppe *et al.* 1986; Ennos *et al.* 1993), and demonstrates that hydrated brace roots have similar geometry between whorls. Lastly, we found that E did not differ between Structural mechanical properties are variable by growth stage within a genotype whorls, however, B73 and A632 brace roots had a higher E than Oh43 (**Supplementary Figure S6**, **Tables S3-4**). There was no apparent relationship between brace root mechanics and the lodging susceptibility. Collectively, these results suggest that brace root mechanics must be interpreted in the context of reproductive stage and genotype to understand how brace root mechanics contribute to root function.

### Structural mechanical properties are variable by growth stage within a genotype

Maize plants are monocarpic and thus the start of reproductive development (R1) marks the end of growth and the start of senescence. Senescence is characterized by the breakdown of cells and remobilization of nutrients, which results in dehydration (Nooden 2012). The loss of turgor pressure accompanied by dehydration has a variable impact on tissue mechanics and depends on whether the cells are thin-walled (e.g. parenchyma) or thick-walled (e.g. sclerenchyma) (Niklas and Spatz 2012). Given the thick-walled structure of maize brace roots (Hoppe *et al.* 1986; Chimungu *et al.* 2015), we hypothesized that dehydration would have a minimal impact on structural mechanical properties. However, when comparing the structural mechanical properties of brace roots between whorls, genotypes, and reproductive stages, we found that the structural mechanical properties of senesced plants are lower than those of hydrated plants (**Figure 6**). There was a significant three-way interaction of whorl by genotype by reproductive stage for K and UL, but the differences were most dramatic for UL (**Table S7-S8**). For UL, whorl 2 was significantly different by stage for all three genotypes, whereas whorl 1 was significantly different by stage for B73 only. As expected, the changes in structural mechanics were not associated with a change in *I* (**Table S7-S8**). In other words, the root geometry does not change during senescence, and thus any changes in structural mechanics are due to the process of senescence itself (e.g., loss of turgor pressure). Collectively, these data suggest that senescence had a significant impact on structural mechanics, and the magnitude of this impact varied by genotype and whorl.

**Figure 6.**
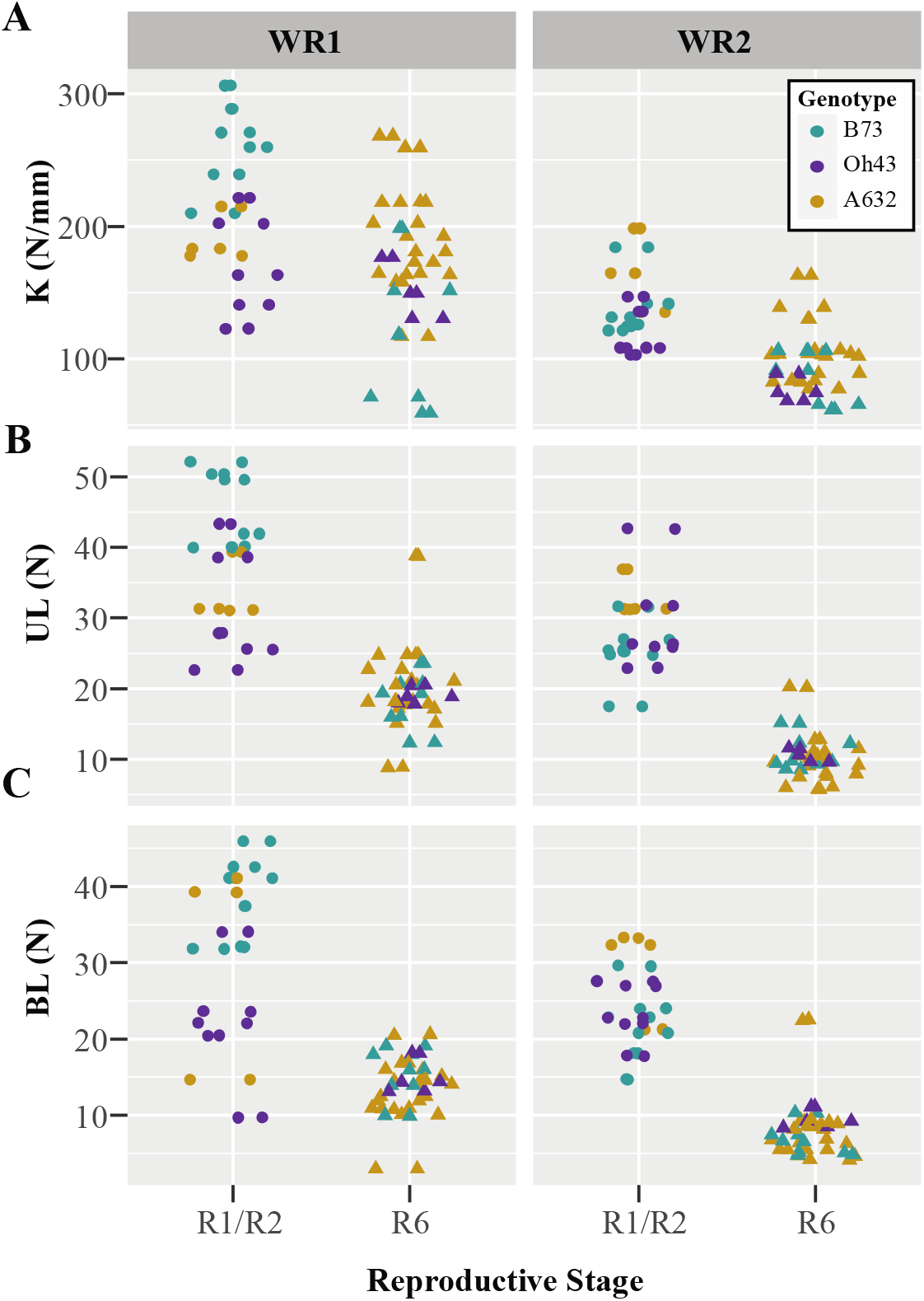
Structural mechanical properties are lower from senesced roots (R6+) compared to hydrated roots (R1/R2). A three-way ANOVA showed that: A) the structural stiffness (K) was impacted by whorl, genotype, and reproductive stage (p≤0.05). Brace roots from the bottom whorl (WR1) of B73 were significantly lower at the R6+ reproductive stage than the R1/R2 reproductive stage. B) The ultimate load (UL) was impacted by whorl, genotype, and reproductive stage (p≤0.05). The UL was significantly lower for the R6+ reproductive stage within the same whorl for B73 (WR1 and WR2), Oh43 (WR2), and A632 (WR2). C) The break load (BL) was not impacted by whorl, genotype, and reproductive stage (p≥0.05). On average brace roots had a lower BL at the R6+ reproductive stage, however, these differences were not statistically significant.

## DISCUSSION

Previous research in maize root biomechanics has shown that nodal roots originating higher on the stalk have progressively higher structural mechanical properties and lower bending moduli (Ennos *et al.* 1993; Goodman and Ennos 2001). However, these previous studies only measured five whorls of crown roots and one whorl of brace roots in one genotype and at one growth stage. Therefore, it was unclear if the same trends in root structural mechanical properties and bending moduli could be extended to additional brace root whorls, genotypes, or growth stages. In this manuscript, we demonstrate that whorl, genotype, and reproductive stage each influence the brace root structural mechanical properties, geometry, and bending moduli.

We have shown that brace roots from whorls closer to the ground contribute more to anchorage, with brace roots from whorl 1 contributing the most (Reneau *et al.* 2020; **Figure 2**). Although roots from whorl 1 were generally stronger than roots from whorl 2, we found a minimal relationship between the brace root contribution ratio of individual whorls and brace root structural mechanical properties (data not shown). This result is consistent with our previous work, which showed that multiple brace root phenotypes are responsible for predicting the brace root contribution to anchorage (Hostetler, Erndwein, *et al.* 2021). Interestingly, among brace root phenotypes, our previous work identified brace root width (i.e., diameter) as the top predictor for the brace root contribution to anchorage at R6+ (Hostetler, Erndwein, et al., 2021). The importance of diameter in this predictive model may be reflective of the importance of brace root structural mechanics, which are partially driven by geometry, as expected, and shown in this study.

When considering differences between brace root whorls within each genotype and reproductive stage, we found that brace roots from higher on the stalk (whorl 2) had lower structural mechanical properties and higher bending moduli compared to brace roots from whorl 1. This finding contradicts previous results from crown roots, which showed that roots from higher nodes have greater strength and lower bending moduli (Ennos *et al.* 1993; Goodman and Ennos 2001). As we have highlighted in this study, the conclusions from crown roots cannot always be extended to brace roots (e.g., increasing diameters). These differences between crown and brace roots are likely driven by the different environments that these roots emerge in - either the soil (crown roots) or air (brace roots). Regardless, these data suggest an inverse relationship between structural mechanical properties and bending moduli within brace root whorls (**Figure 3, Figure 5**). One explanation for this inverse relationship is that larger diameter roots have higher structural mechanics since they can distribute loads over a greater area but have relatively less structural tissues. In other words, maize roots have been considered as two concentric structural cylinders of lignified tissue, with non-structural intervening parenchyma tissues (Ennos *et al.* 1993; Chimungu *et al.* 2015). Under this assumption, roots with a larger diameter root have more parenchymal area, which leads to lower overall material mechanics. Our results support this assumption, and emphasize the importance for future studies to consider the whorl-origin of the nodal roots and whether this node is located above or below the ground.

Our results further expand on previous studies in one genotype to demonstrate that there is variation in brace root structural mechanical properties and bending moduli between genotypes. Interestingly, genotypes with higher structural mechanical properties also have higher bending moduli at both reproductive stages. Thus, suggesting that the inverse relationship between structural mechanical properties and bending moduli observed between whorls is a within-genotype phenomenon. Overall, these results highlight the potential for future work aimed at determining the genetic and environmental regulation of brace root mechanics for crop improvement.

Lastly, we aimed to understand how senescence impacts brace root mechanics. We posited that there would be minimal changes in structural mechanical properties between reproductive stages, because brace roots are composed of thick-walled structural elements (Hoppe *et al.* 1986; Chimungu *et al.* 2015), which are less impacted by turgor pressure (Niklas and Spatz 2012). However, our results show that the structural mechanical properties are lower after senescence, suggesting that the contribution of thin-walled parenchyma cells is impacting the overall mechanics. This suggests that the concentric rings of lignified tissues are not the only tissues contributing significantly to brace root mechanics. Future work aimed at dissecting the tissue-specific contribution of each cell layer to the overall structural mechanics will be important to refine targets for crop improvement.

The data presented here expand our understanding of the factors that impact maize root mechanics and show variation in brace root mechanics is specific to whorl-origin, genotype, and reproductive stage. Interestingly, despite the variation we observed in this study, the bending moduli are within range of previous reports on root mechanics (**Figure 7**). In contrast, the bending moduli of senesced stalks are much greater, suggesting that stalks can resist bending more than roots. One explanation for the different bending moduli ranges is that roots must retain the ability to flex and absorb forces, which is likely a key strategy to maintain anchorage (Stubbs *et al.* 2019). Overall, this work sets the foundation to address additional open questions about the genetic and environmental basis of root mechanics, the functional consequences of mechanical variation, and the underlying tissue mechanics that lead to organ-level structural mechanics.

**Figure 7.**
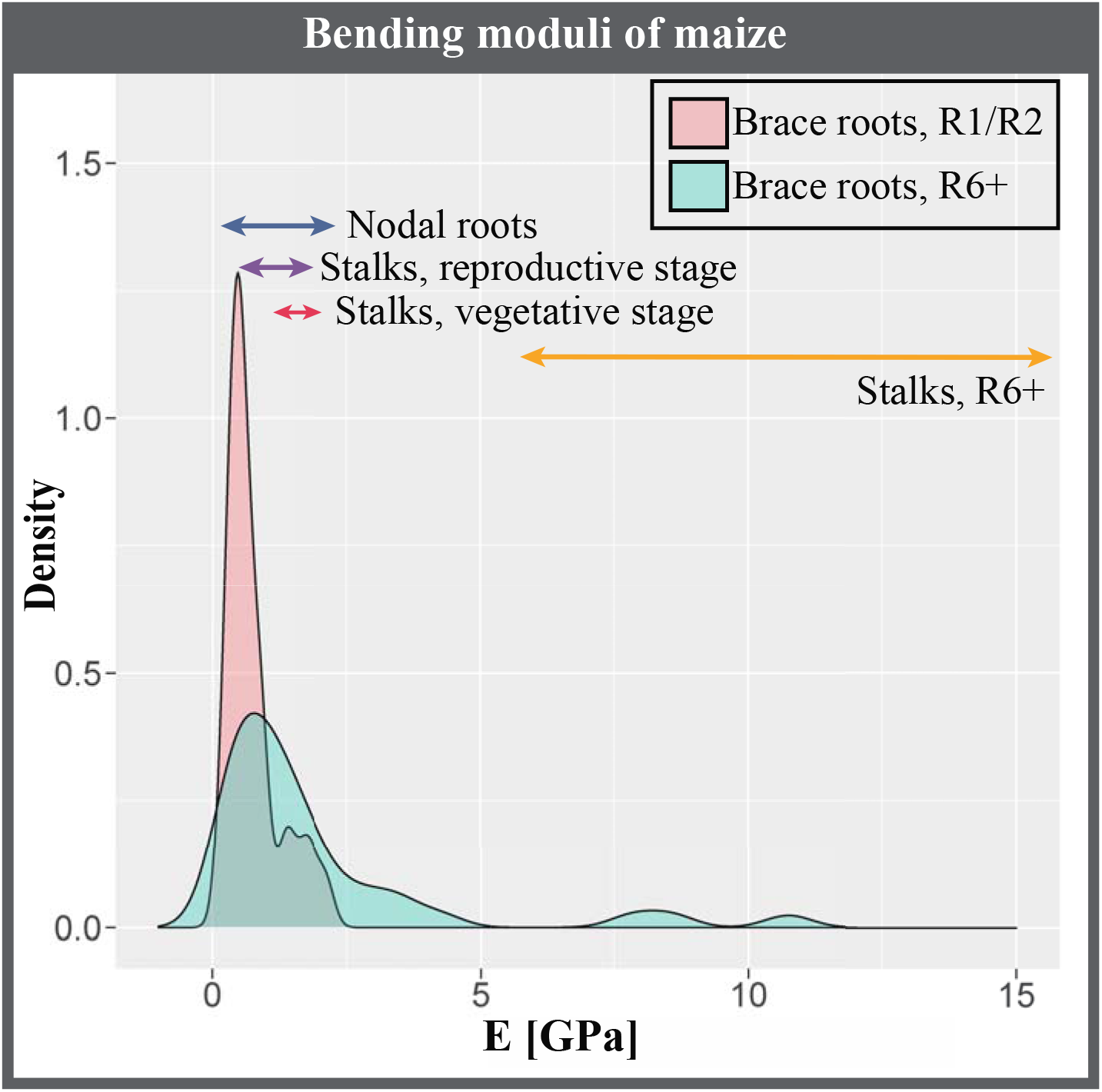
The bending moduli of maize brace roots at the R6+ and R1/R2 reproductive stage are like those of other studies. A comparison of the bending moduli of the brace roots from this study with the bending moduli of maize roots and stems from previous studies shows that brace root samples are comparable with other maize roots samples. Nodal roots data is from (Ennos *et al.* 1993; Goodman and Ennos 2001); Stalks, reproductive stage data is from (Goodman and Ennos 1997, 1998); Stalks, vegetative stage data is from (Goodman and Ennos 1996); stalks, R6+ data is from (Al-Zube *et al.* 2018)

## ACKNOWLEDGEMENTS

We gratefully acknowledge the members of the Sparks lab for assistance in harvesting plants, collecting brace roots, and preparing brace root samples. We additionally acknowledge Rich West and the Center for Biomedical and Brain Imaging (CBBI) for microCT assistance, Dr. Dawn Elliot (University of Delaware) for access to Instron, Dr. Douglas Cook (Brigham Young University) for assistance identifying foam support for microCT scanning, and Dr. Rubén Rellán-Álvarez (University of North Carolina) for providing the picture of root lodging. This research was made possible by funding from the University of Delaware Research Foundation and the Thomas Jefferson Fund to EES.

## AUTHOR CONTRIBUTIONS

LE, EG, MLK, and EES conceptualized the project. LE, EG, JWR, and EES collected data. ANH, LE, EG, MLK, and EES analyzed data. All authors contributed to the writing and/or editing of the manuscript.

## SUPPLEMENTARY DATA

**Figure S1.**
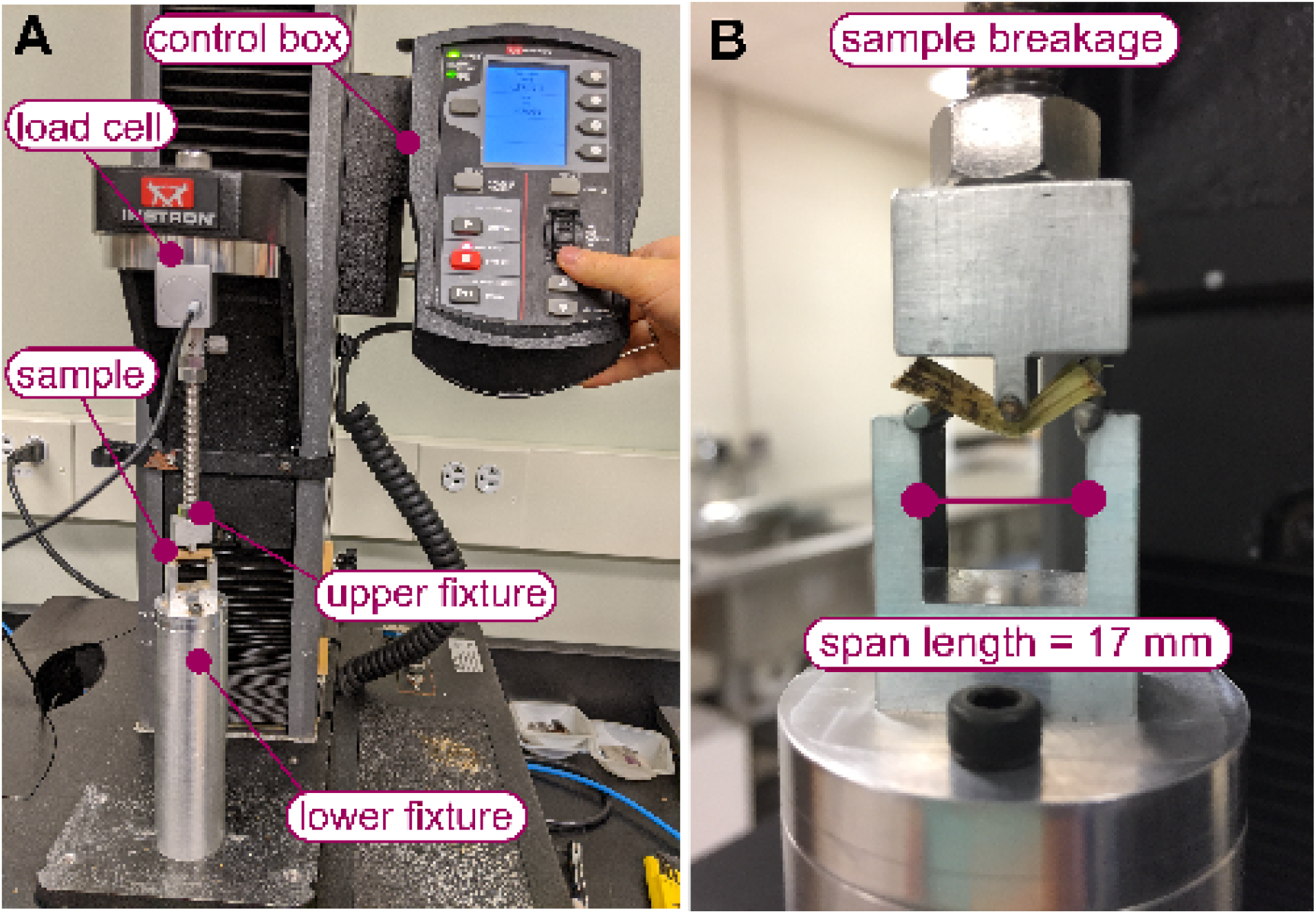
Overview of the setup for 3-point bend testing. A) Brace root samples were tested using an Instron 5943 (Norwood, Massachusetts USA) equipped with a 100 N load cell (Instron 2530 Series static load cell, Norwood, Massachusetts USA). Samples were placed on a lower fixture, which supported the sample at both ends. An upper fixture attached to a 100 N load cell was moved vertically in contact with the center of the sample. Each sample was preloaded to 0.2 N and the displacement readout was calibrated. Each test proceeded at a rate of 1 mm/min and lasted about 90 seconds each (until failure was detected). B) Brace roots were closely monitored during testing to ensure that failure occurred on the sample surface opposite the anvil.

**Figure S2.**
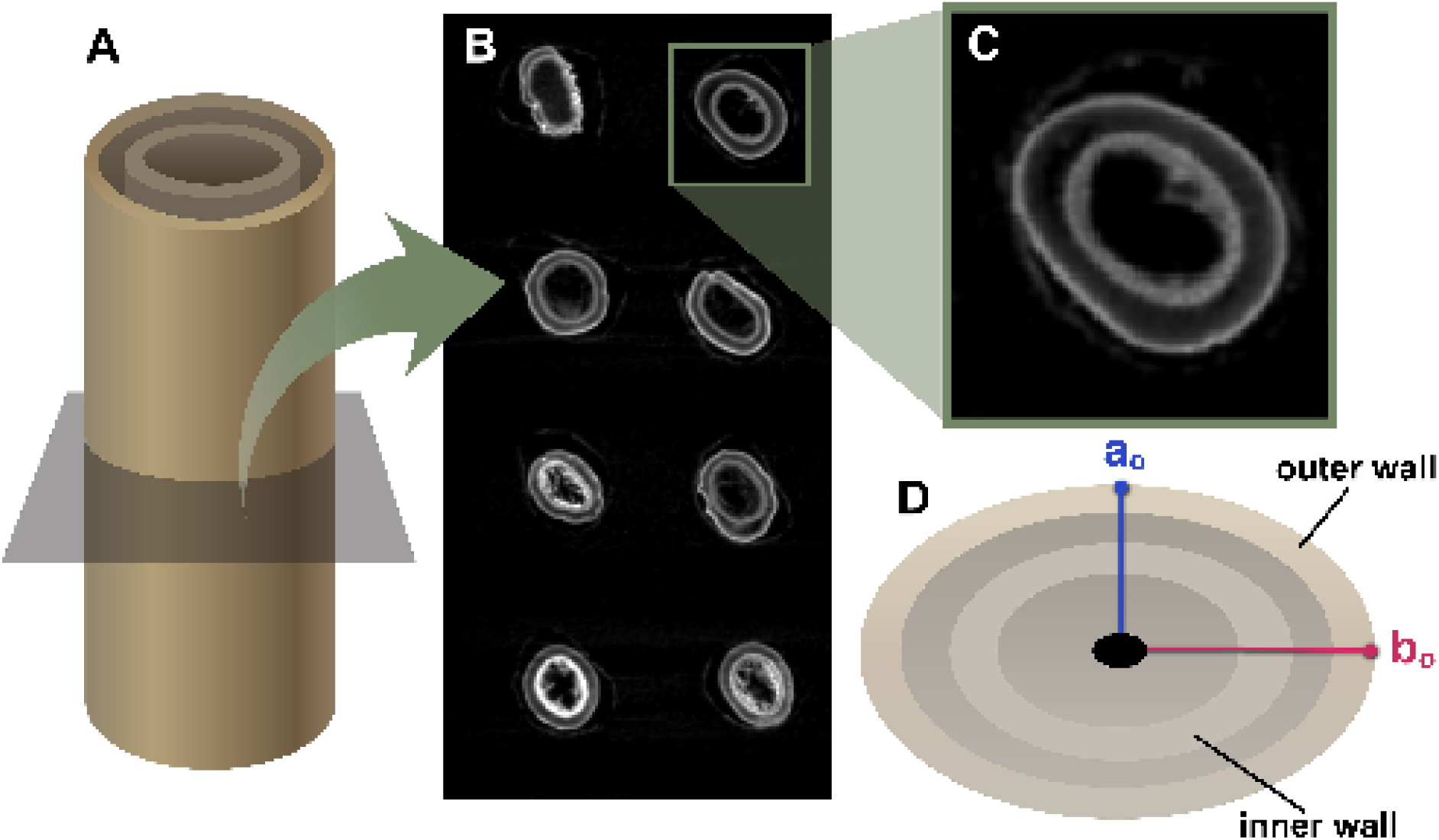
Quantification of R6+ brace root geometry from microCT scans. Multiple brace root samples were loaded into a custom bed made of low-density upholstery foam for microCT scanning. A) R6+ brace root geometries were measured using the central cross section microCT images (the location in contact with the upper anvil of the mechanical testing fixture). B) Brace root samples were inserted into a low-density upholstery foam fixture, which provided a supportive bed and did not appear on the microCT scan image. C) An example microCT scan illustrates the hollow double-walled geometry of brace root cross sections. D) The following dimensions were measured from the central cross section of microCT scans: Distance from the center point to the exterior of the minor outer wall perpendicular to bending (ao), and to the exterior of the major outer wall parallel to bending (b_o_).

**Figure S3.**
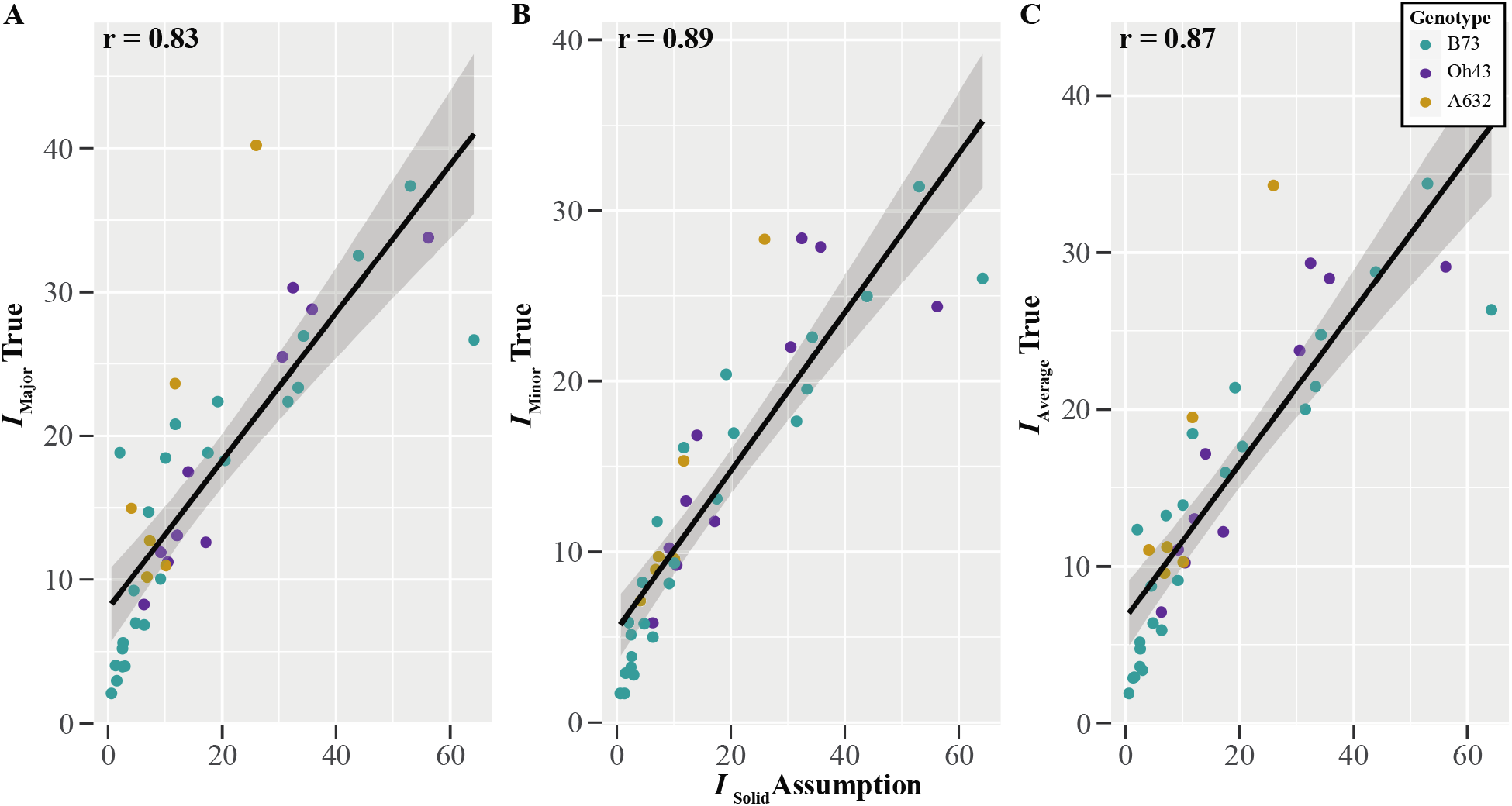
The simplified assumption of the second moment of area (*I*) is a reasonable approximation of the true *I*. The second moment of area was calculated directly from the microCT images (*I*_true_ major, *I*_true_ minor, and *I*_true_ average) and compared with *I* calculated from a simplified solid cylinder assumption. A Pearson correlation analysis showed that A) *I*_solid_ was positively correlated with A) *I*_true_ major with an r = 0.83, B) *I*_true_ minor with an r = 0.89, and C) *I*_true_ average with an r=0.87.

**Figure S4.**
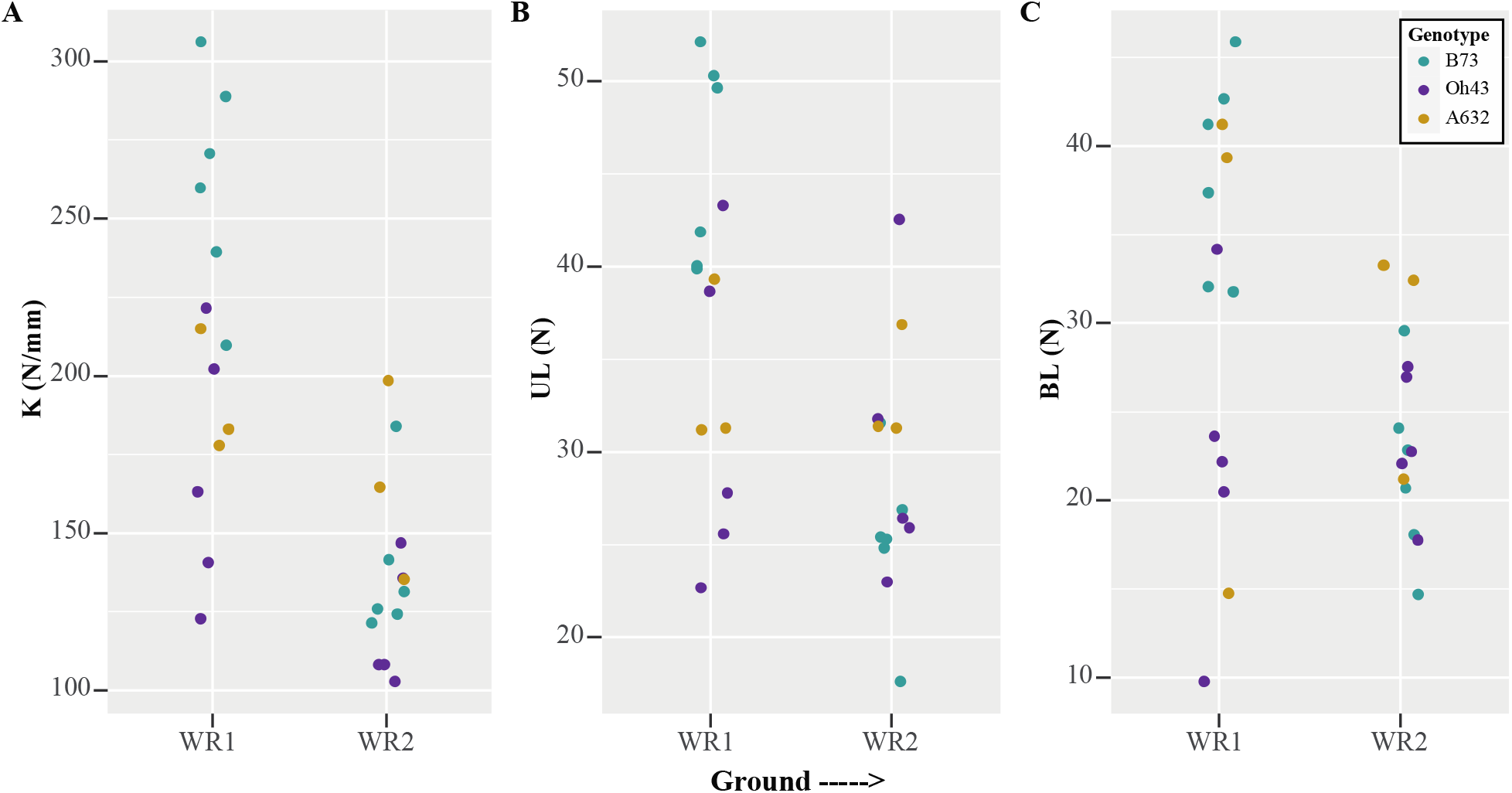
The variation in structural mechanical properties of maize brace roots depends on genotype for the R1/R2 reproductive stage. A) A two-way ANOVA showed that the variation in structural stiffness (K) of maize brace roots between whorls was dependent on genotype (p≥0.05) at the R1/R2 reproductive stage. The K was higher for whorl 1 compared to whorl 2 for B73 and Oh43 (p≥0.05), however whorls did not differ in K for A632 (p≥0.05). B) A two-way ANOVA showed that the variation in the ultimate load (UL) of maize brace roots between whorls was dependent on genotype (p≥0.05), with B73 being the only genotype that had a statistically higher UL for whorl 1 compared to whorl 2. C) A two-way ANOVA showed that the variation in the break load (BL) of maize brace root whorls was dependent on genotype (p≥0.05), with B73 being the only genotype that had a statistically higher BL for whorl 1 compared to whorl 2. WR – whorl.

**Figure S5.**
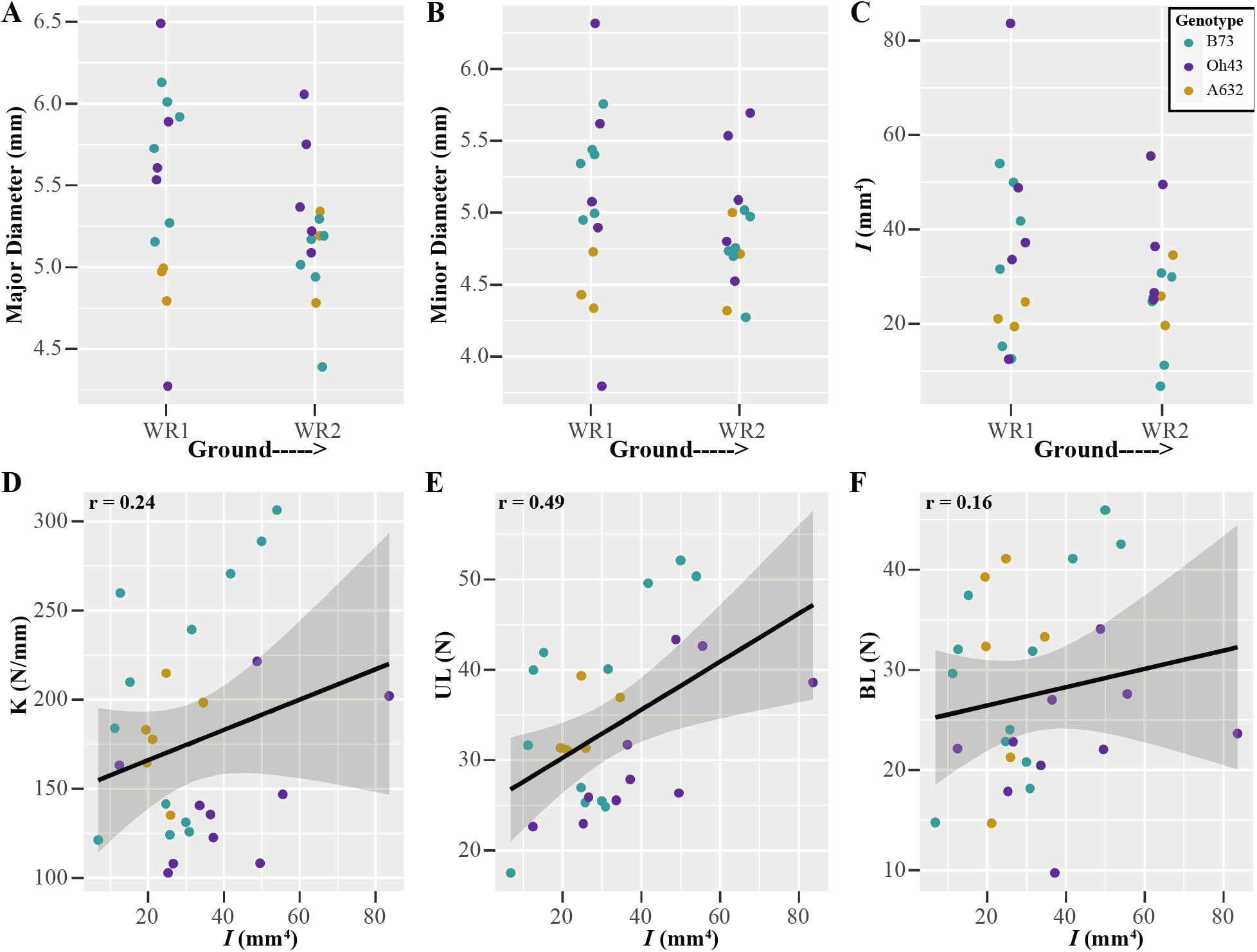
Brace root geometry does not differ among whorls or genotypes at the R1/R2 reproductive stage. A two-way ANOVA showed that the A) major diameter, B) minor diameter, and C) second moment of area (*I*) did not differ among whorls or genotypes at the R1/R2 reproductive stage (p≥0.05). WR – whorl.

**Figure S6.**
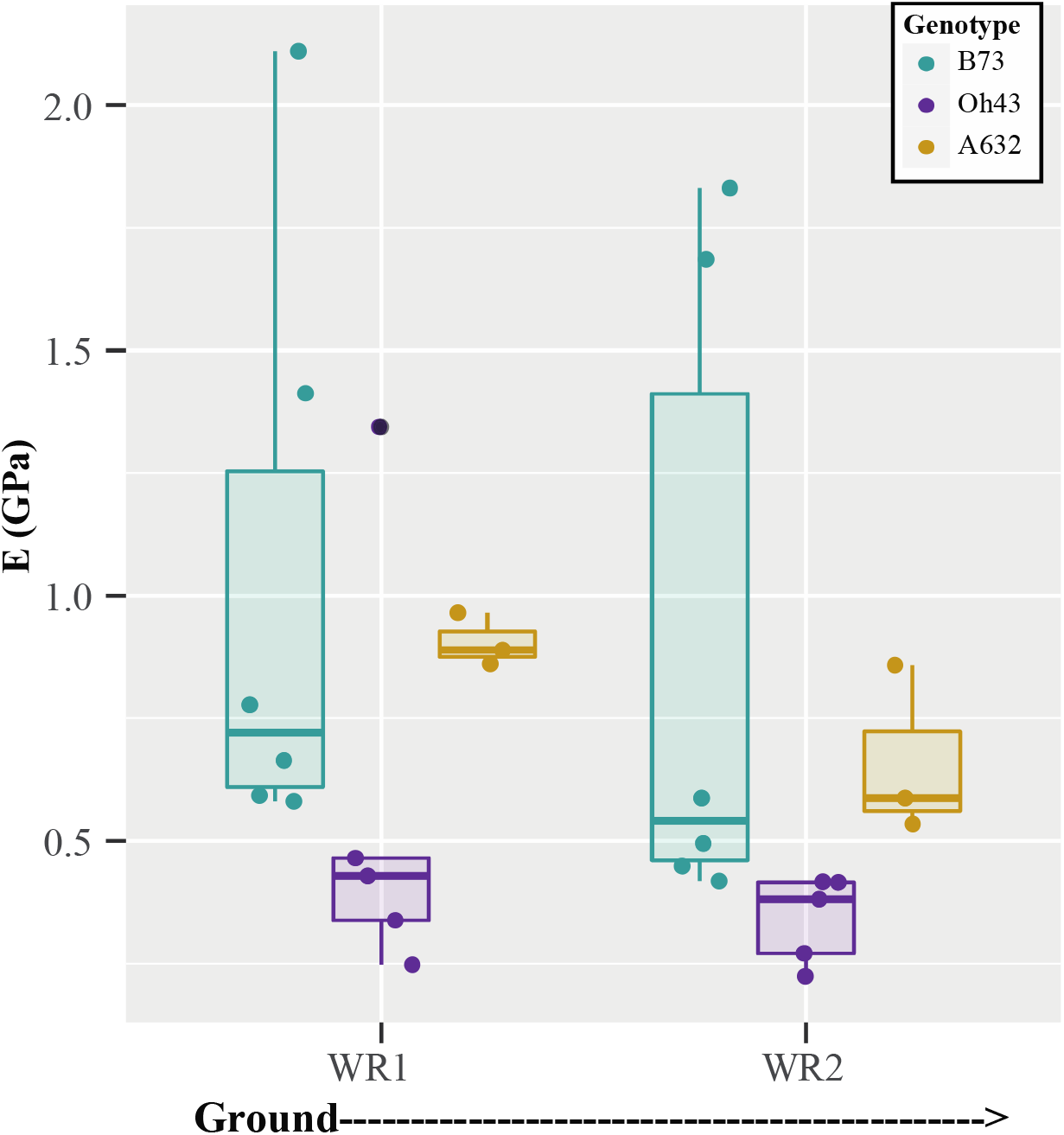
Brace root material properties vary by genotype but not whorl for the R1/R2 reproductive stage. A two-way ANOVA showed that the bending modulus (E) of maize brace roots did not differ by whorl (p≥0.05) at the R1/R2 reproductive stage. However, E did differ between genotypes (p≥0.05), with B73 and A632 having significantly higher material properties compared to Oh43. WR – whorl.

**Table S1. Two-way ANOVA results for the brace root contribution to anchorage ratio and the ratio of individual whorls.** A two-way ANOVA was run for the brace root contribution to anchorage ratio (<u>b</u>race <u>r</u>oot <u>c</u>ontribution <u>r</u>atio, BRCR) with the number of whorls in the ground and genotype as the independent variables. Both the number of whorls in the ground and genotype were significant (p≥0.05), but there was no interaction between the two. A separate two-way ANOVA was run for the BRCR for individual whorls with whorl and genotype as the independent variables. Both whorl and genotype were significant (p≥0.05), but there was no interaction between the two. The gray divider illustrates individual ANOVA tables.

**Table S2. Pairwise comparisons for the brace root contribution to anchorage ratio and the ratio of individual whorls.** A post-hoc Tukey’s Honest Significant Difference (HSD) test was used to determine genotypes that were significantly different for the brace root contribution to anchorage ratio (<u>b</u>race <u>r</u>oot <u>c</u>ontribution <u>r</u>atio, BRCR) and the BRCR for individual whorls. Lowercase letters indicate when whorls were significantly different from one another, whereas uppercase letters indicate when genotypes were significantly different from one another. Groups that share a letter (case matters) indicate no significant difference (p≥0.05). Data are group means ± standard deviation (SD). A minimum of two roots per plant and three plants per genotype were used to calculate group means. The gray divider illustrates individual TukeyHSD results.

**Table S3. Two-way ANOVA results for the structural mechanical properties, geometry, and material properties of maize brace roots at the R6+ and R1/R2 reproductive stage.** A two-way ANOVA was run for individual data sets (K, UL, BL, MajorD, MinorD, *I*, and E) at each reproductive stage (R6+ and R1/R2) individually. The gray divider illustrates individual ANOVA tables. For each ANOVA, whorl and genotype were the independent variables. For the R6+ data, differences were found between whorls (p≥0.05) for structural mechanical properties, geometry, and material properties. Additionally, a genotype effect (p≥0.05) was found for specific traits (K, MinorD, *I*, and E). For the R1/R2 data, differences in structural mechanical properties were dependent on genotype (p≥0.05 for the interaction). There were no differences found for measures of geometry, and the bending modulus differed between genotypes (p≥0.05) but not whorls (p≥0.05).

**Table S4. Pairwise comparisons for the structural mechanical properties and material properties of maize brace roots at the R6+ and R1/R2 reproductive stage.** A post-hoc Tukey’s Honest Significant Difference (HSD) test was used to identify differences in whorls and genotypes for structural mechanical properties, and differences in genotypes for material properties. Lowercase letters indicate when whorls and genotypes were significantly different from one another for structural mechanical properties, whereas uppercase letters indicate when genotypes were significantly different from one another for material properties. Groups that share a letter (case matters) indicate no significant difference (p≥0.05). Data are group means ± standard deviation (SD). A minimum of two roots per plant and three plants per genotype were used to calculate group means. The gray divider illustrates individual TukeyHSD results.

**Table S5. Two-way ANOVA results for the bending modulus calculated from *I*_solid, true_ major, *I*_true_ minor, and *I*_true_ average.** Regardless of how the bending modulus was calculated (from *I*_solid_, *I*_true_ major, *I*_true_ minor, or *I*_true_ average), whorl and genotype had the same effect. A two-way ANOVA showed that whorl (p≥0.05) and genotype (p≥0.05) impacted E. The gray divider illustrates individual ANOVA tables.

**Table S6. Pairwise comparisons for the bending modulus calculated from *I*_true_ major, *I*_true_ minor, and *I*_true_ average.** A post-hoc Tukey’s Honest Significant Difference (HSD) test was used to identify differences in whorls and genotypes for the different calculations of E (either calculated from *I*_true_ major, *I*_true_ minor, or *I*_true_ average). Lowercase letters indicate when whorls were significantly different from one another for E, whereas uppercase letters indicate when genotypes were significantly different from one another for E. Groups that share a letter (case matters) indicate no significant difference (p≥0.05). Data are group means ± standard deviation (SD). A minimum of 2 roots per plant and three plants per genotype were used to calculate group means. The gray divider illustrates individual TukeyHSD results.

**Table S7. Three-way ANOVA results for the structural mechanical properties, geometry, and the second moment of area (*I*) of maize brace roots.** A three-way ANOVA was run to determine the effect of reproductive stage (R1/R2 or R6+), whorl, and genotype on structural mechanical properties and geometry (*I*). For the structural stiffness (K) and ultimate load (UL), a three-way interaction was observed (p≥0.05). For break load (BL) and *I*, a three-way interaction was not observed (p≥0.05). The gray divider illustrates individual ANOVA tables.

**Table S8. Pairwise comparisons for the structural mechanical properties and the second moment of area (*I*) of maize brace roots.** A post-hoc Tukey’s Honest Significant Difference (HSD) test was used to identify differences in reproductive stages for structural stiffness (K) and the ultimate load (UL). Groups that share a lowercase letter indicate that a significant difference was not observed (p≥0.05). Data are group means ± standard deviation. A minimum of two roots per plant and three plants per genotype were used to calculate group means. The gray divider illustrates individual TukeyHSD results.

